# The taxonomic status of *Farlowella colombiensis* Retzer & Page 1997 with comments on species of the *F. acus* species group (Loricariidae: Loricariinae)

**DOI:** 10.1101/2024.11.18.624099

**Authors:** Omar Eduardo Melo-Ortiz, Michael E. Retzer, Saúl Prada-Pedreros, Gustavo A. Ballen

**Author notes:** Corresponding author: Gustavo A. Ballen. Instituto de Biociências, Universidade Estadual Paulista “Júlio Mesquita Filho”, Botucatu, SP, Brazil. **Funding statement:** This work was funded by FAPESP processes 2014/11558-5, 2016/02253-1, and 2023/07838-1 to GAB.

## Abstract

The genus *Farlowella* has been historically challenging, in part due to the difficulty in defining diagnostic characters which allow to clearly set apart and identify the species. *Farlowella colombiensis* Retzer & Page 1997 is one of such examples, whose diagnostic characters were based on caudal-fin colour pattern, body cover ventral pattern, and details of the head. We herein reassess the taxonomic status of this species against congeners of the *F. acus* species group (*F. acus*, *F. martini*, *F. mitoupibo*, *F. venezuelensis*, and *F. vittata*). We found no significant differences between *F. colombiensis* and *F. acus* in morphometric, meristic, and discrete characters, therefore rendering *Farlowella colombiensis* a junior synonym of *Farlowella acus*. We provide remarks on different species of the *F. acus* group in Colombia, as well as description of sexual dimorphism in the genital papilla for the first time in the subfamily. We also provide a key to the species of the *F. acus* species group in Colombia.

## Introduction

The genus *Farlowella* Eigenmann & Eigenmann (1889) is the second richest genus in the subfamily Loricariinae (Terán et al., 2019; Dopazo et al., 2023), with 32 valid species (Fricke et al., 2024). At least 13 species are found in Colombia (DoNascimiento et al 2024), mostly in drainages east of the Andes (i.e., Orinoco and Amazonas), and marginally in the Maracaibo and Magdalena-Cauca basins (Ballen & Mojica, 2014; Retzer & Page, 1997). *Farlowella* is readily diagnosable because of its cylindrical body, which is also elongate and gracile, with a very produced snout of variable length, and dorsal fin inserted opposed to the anal fin (Boeseman, 1971; Burgess, 1989; Covain & Fisch-Muller, 2007; Eigenmann & Vance, 1917; Kner, 1853; Regan, 1904; Retzer & Page, 1997).

Although the genus is well-defined using external morphology, its taxonomy is still challenging and some studies have found it to be paraphyletic with respect to the monotypic *Aposturisoma* (Covain et al., 2016). Retzer & Page (1997) revised the genus for the first time recognising 24 species, six of them newly described. They arranged the genus in six species groups: *F. acus*, *F. amazona*, *F. curtirostra*, *F. knerii*, *F. mariaelenae,* and *F. nattereri*. However, some species of uncertain relationships remained unassigned to a group (*F. gracillis*, *F. hahni*, *F. oxyrryncha*, *F. paraguayensis*, *F. reticulata*, and *F. smithi*).

Some species have been described after Retzer & Page’s revision, mostly without changes to the definition of the species groups (e.g., *F. altocorpus*, *F. azpelicuetae*, *F. gianetii*, *F. guarani*, and *F. azpelicuetae*, assigned to the *F. nattereri* group, and *F. wuyjugu*, unassigned to group; Ballen et al., 2016a; Retzer, 2006; Terán et al., 2019, Delgadillo et al., 2021; Dopazo et al., 2023) whereas species descriptions have sometimes resulted in species groups having been found to be less defined (e.g., *F. mitoupibo* and *F. yarigui*, assigned to the *F. acus* group; Ballen & Mojica, 2014; Ballen et al., 2016b).

Despite efforts towards understanding the taxonomy of the genus, there is still ongoing uncertainty on the status of several species, and additional ones being described lately (e.g., Dopazo et al., 2023). Part of the issue lies in the presence of variation for diagnosing some groups or species (e.g., Ballen et al., 2016b; Dopazo et al., 2023), whereas in other cases the lack of obvious external features make it challenging to distinguish the species. One example of this variation is found in *F. colombiensis*, which has two complete abdominal rows, whereas some individuals present a third, median, incomplete one, which has been proposed as a diagnostic character for the species (Retzer & Page, 1997). Such a character has been also found in *F*. *mitoupibo*, another member of the *F. acus* group (Ballen et al., 2016b). This has casted doubts on whether such characters are useful for diagnosing the species group from others in the genus.

Retzer & Page (1997) suggested that *F. colombiensis* can be distinguished from congeners using morphometric characters, however, these show overlap with *F. acus*. So far no additional characters have been suggested for diagnosing the species besides the presence of a middorsal crest on the head which is also present in other species of the *F. acus* group (e.g., *F. martini* and *F. venezuelensis*).

The main goal of this paper is to assess the taxonomic status of *Farlowella colombiensis* and to provide a redescription for *Farlowella acus*, with comments on the congeneric members of the *F. acus* group in Colombia. Furthermore, we present for the first time evidence of sexual dimorphism in the urogenital papilla of the genus *Farlowella*.

## Materials and methods

Terminology, meristics, and morphometrics follow Ballen et al. (2016b), with the addition of distance between the posterior margin of nare and anterior margin of the eye (eye-nare distance). Plate terminology follows Ballen et al. (2016a). Meristics are reported with frequency between parentheses. Line morphometric variables were taken with Mitutoyo digital callipers of precision 0.01 mm. Standard length is in millimetres (mm), whereas remaining measurements are proportions of Ls, except for subunits of the head, where proportions are presented with respect to the head length.

A total of 355 individuos were examined at the Ichthyology Collection, Museo Javeriano de Historia Natural "Lorenzo Uribe, S. J." (MPUJ).

Sexual dimorphism in mature individuals was determined by the presence of nuptial odontodes on the sides of the snout in males. Females were identified due to a bulky venter and absence of nuptial odontodes on the snout. We identified herein dimorphism in the urogenital papilla for the first time in the genus, which allows to effectively separate mature males from females.

### Statistical analyses

We analysed the quantitative data using R v.4.4.1 (R Core Development Team, 2024), available at http://www.r-project.org. We used a Principal component analysis (PCA) in order to reduce dimensionality and explore potential information in our multivariate dataset, using the packages FactoMineR and factoextra (Lê et al., 2008; Kassambara and Mundt, 2020) scaling to unit dimensions with the argument "scale.unit = TRUE" in the function FactoMineR::PCA, which avoids overweight by differences in order of magnitude among the variables in the dataset. We chose to analyse proportions of each variable against either Ls or head length in order to remove the size effect which would otherwise be present in the principal component 1. Scatterplots were generated with the package car (Fox & Weisberg, 2019). Nonparametric statistics were implemented with the package npsm and following Kloke & McKean (2014 a, b). We explored possible differences among species as well as between sexes. We used the body width at the insertion of the dorsal fin as a covariate to snout measurements in order to guarantee independence between variables, avoiding possible general scalings which affect the head measurements as a whole. We emphasised comparisons between putative specimens of *F. acus* and *F. colombiensis*, even though our morphometric dataset included all the species of the *F. acus* group, because our present focus is the taxonomic status of *F. colombiensis*.

## Results

We identified four species assigned to the *F. acus* group among the material examined from Colombia: *F. vittata* (Figure 1b-c), *F*. *acus* (Figure 1d-f), *F*. *colombiensis* (Figure 1g-h), and *F*. *mitoupibo* (Figure 1i). Morphometric variables are presented in Table 1, and meristics in Table 2.

**Figure 1.**
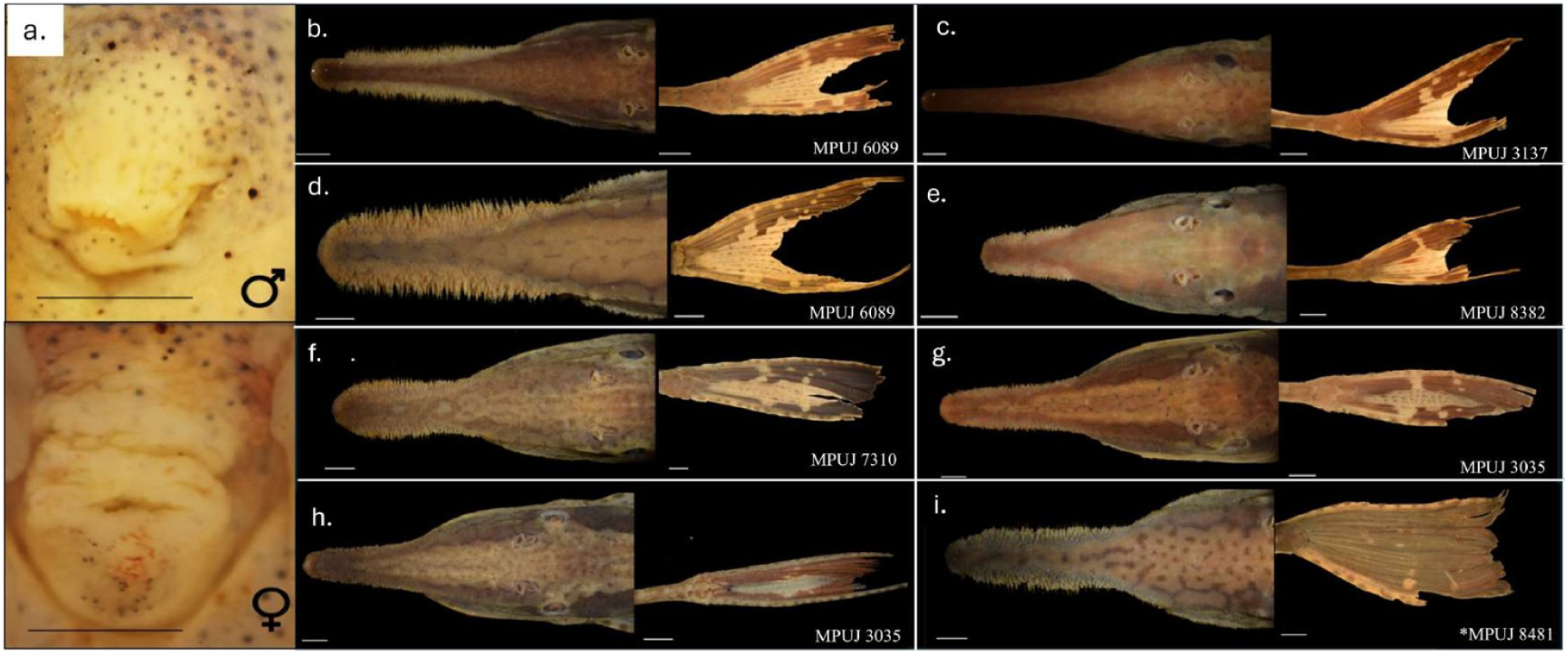
Morphological variation within species of the genus *Farlowella* belonging to the *F. acus* group. a. Variation of the urogenital papilla, upper is male, lower is female; b – c. *F. vittata* (MPUJ 6089 and 3137); d – f. *F. acus* (MPUJ, 6089, 8382, 7310); g – h. *F. colombiensis* (MPUJ 3035); and i. *F. mitoupibo* (MPUJ 8481, Holotype). The black line represents a 2mm scale; the white line represents a 3mm scale.

**Table 1.**
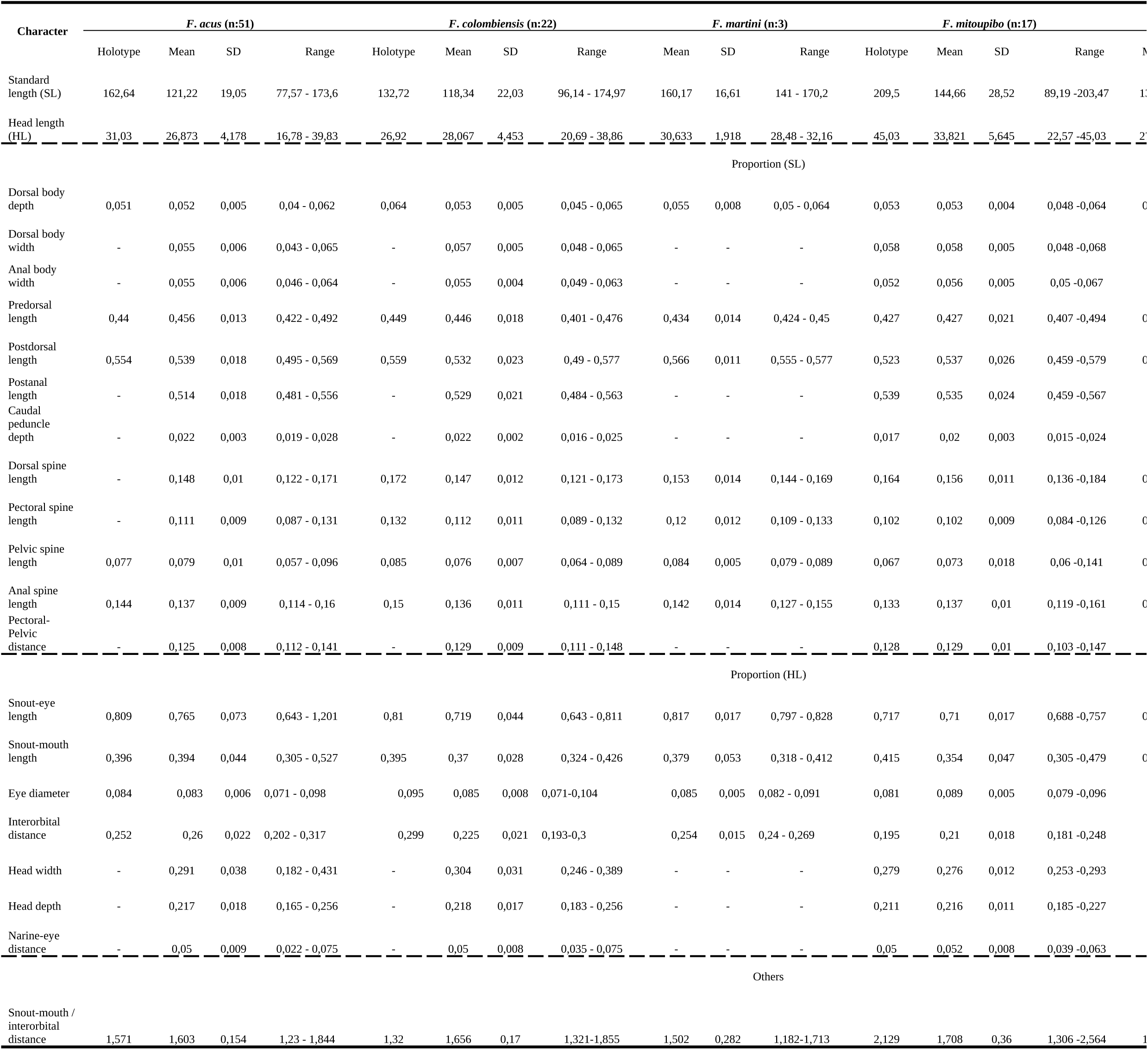
Morphometric data of median, standard deviation (SD), and range for *F. acus*, *F. colombiensis*, *F. martini*, *F. mitoupibo*, *F. venezuelensis*, and *F. vittata*. The range for *F. acus*, *F. colombiensis* and *F. mitoupibo*, includes the holotype (NMW 47795, CAS-SU (ICH) 23733 and MPUJ 8481, respectively).

**Table 2.**
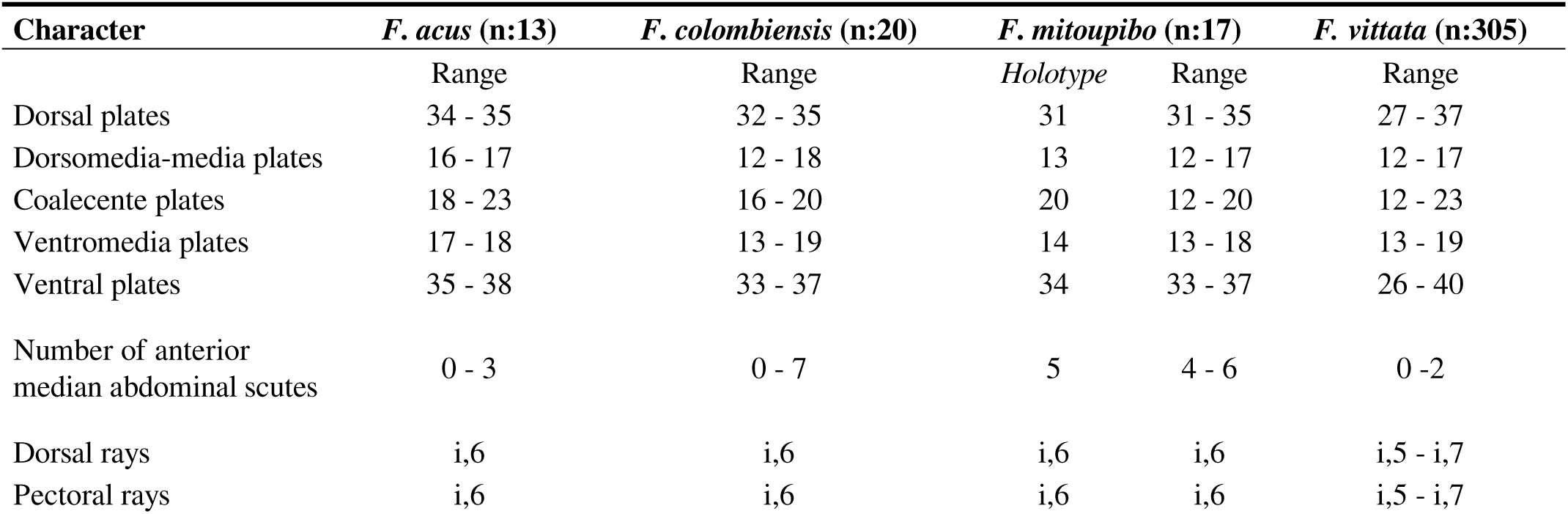
Meristic counts of *F. acus*, *F. “colombiensis”*, *F. mitoupibo*, and *F. vittata*. The range for *F. acus*, *F. “colombiensis”* and *F. mitoupibo* includes the holotype (NMW 47795, CAS-SU (ICH) 23733 and MPUJ 8481, respectively).

The PCA (Figure 2) showed overlap between *F. mitoupibo* and the rest of species, as well as in *F. vittata*. However, this overlap was complete between specimens of *F. acus, F*. *colombiensis*, *F. martini* and *F. venezuelensis*. At the moment, the only way to distinguish these species is using the geographic distribution, as *F. martini* and *F. venezuelensis* are restricted to Venezuela (Figure 3a-b). We defer any further comment on the Venezuelan species as our sampling was poor for that country, and emphasise the comparisons between *F. acus* and *F. colombiensis* from now on, because of the overlapping distribution (Figure 3c-d).

**Figure 2.**
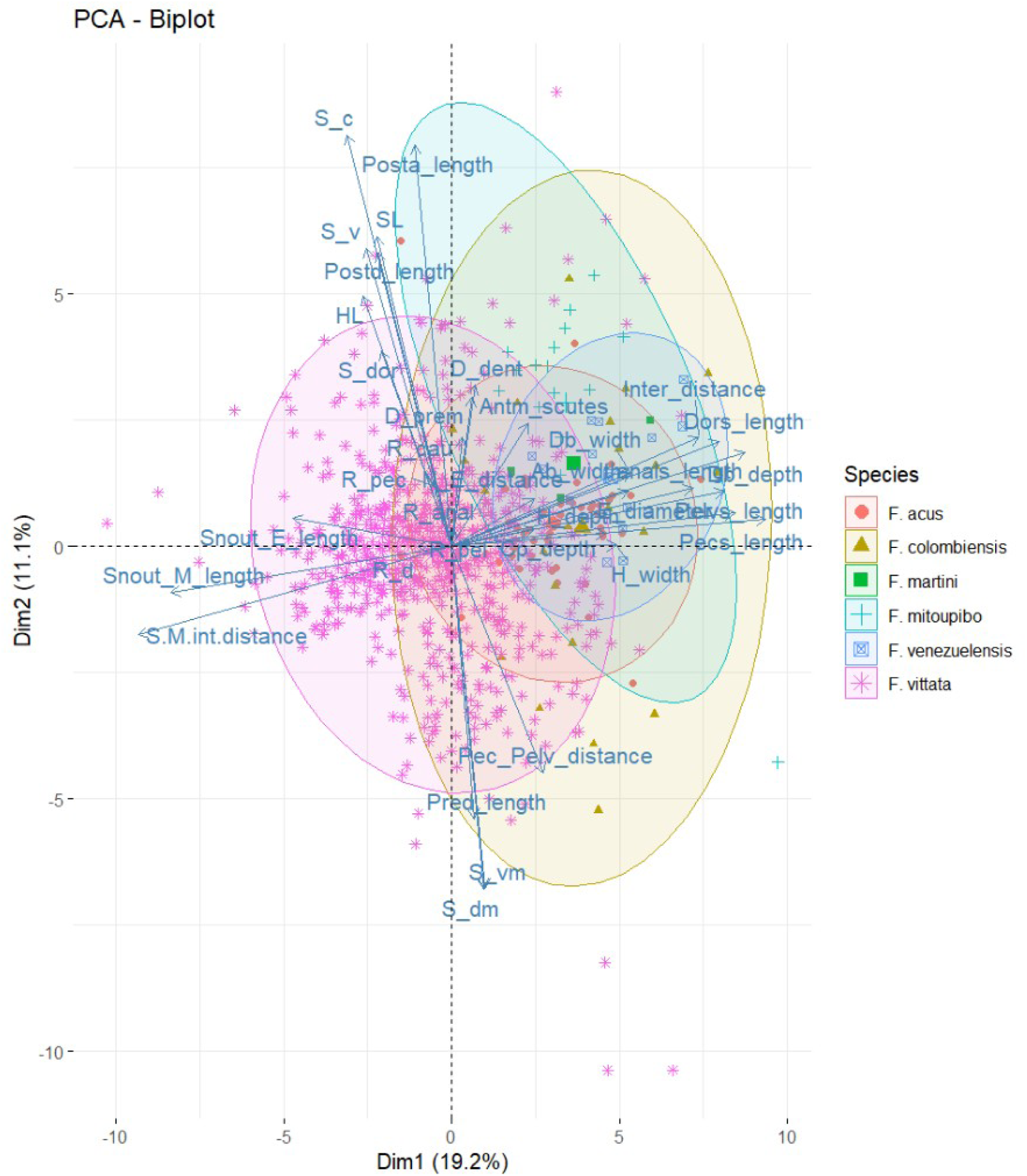
Principal component analysis (PCA) for the morphometric and meristic variables among species of the *Farlowella acus* species group: *F. acus*, *F. colombiensis*, *F. martini*, *F. venezuelensis*, *F. vittata* and *F. mitoupibo*. The biplot depicting PC1 and PC2, and explains 30.3% of the variance. SL:Standard length, HL:Head length, Db_depth:Dorsal body depth, Db_width:Dorsal body width, Ab_width:Anal body width, Pred_length:Predorsal length, Postd_length:Postdorsal length, Posta_length:Postanal length, Cp_depth:Caudal peduncle depth, Dors_length:Dorsal spine length, Pecs_length:Pectoral spine length, Pelvs_length:Pelvic spine length, Anals_length:Anal spine length, Pec_Pelv_distance:Pectoral-Pelvic distance, Snout_E_length:Snout-eye length, Snout_M_length:Snout-mouth length, E_diameter:Eye diameter, Inter_distance:Interorbital distance, H_width:Head width, H_depth:Head depth, N_E_distance:Narine-eye distance, S-M/int-distance:Snout-mouth / interorbital distance, S_dor:Dorsal plates, S_dm:Dorsomedia-media plates, S_c:Coalecente plates, S_vm:Ventromedia plates, S_v:Ventral plates, R_d:Dorsal rays, R_pec:Pectoral rays, R_pel:Pelvic rays, R_anal:Anal rays, R_cau:Caudal rays, D_prem:Premaxillary teeth, D_dent:Dentary teeth, Antm_scutes:Number of anterior median amdominal scutes.

**Figure 3.**
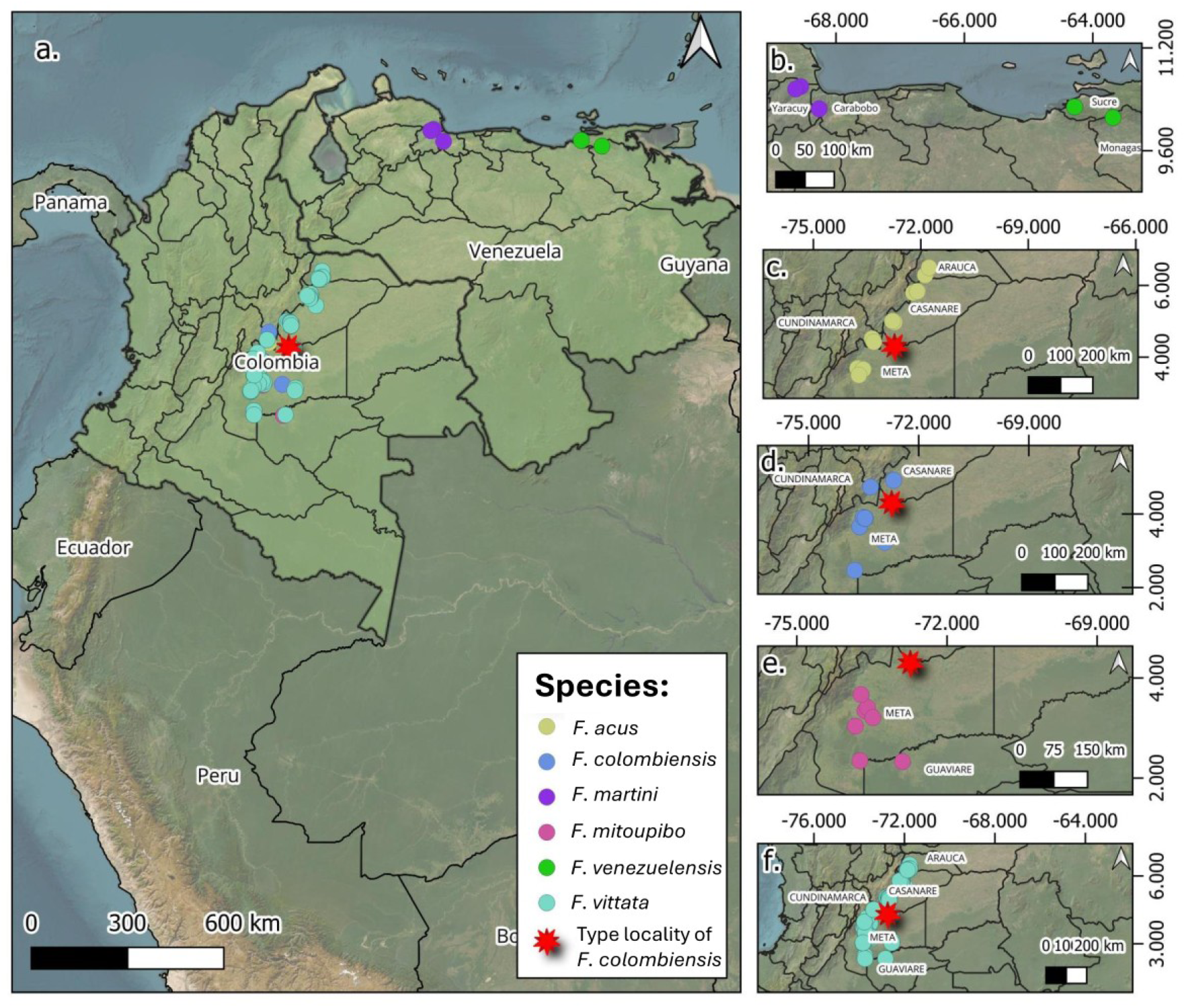
Geographic distribution of species in the *F. acus* group. a. Overview of the distribution in Colombia and Venezuela; b. *F. martini* and *F. venezuelensis*; c. *F. acus*; d. *F. colombiensis*; e. *F. mitoupibo*; f. *F*. *vittata*. The star represents the type locality of *Farlowella colombiensis* (Río Upia).

After examining multiple specimens assigned to *F. acus* and *F. colombiensis*, we conclude that they are synonyms, as supported by the PCA, ANCOVA, and testing of diagnostic characters (Figures 1d-h,4). We present below a redescription for *F. acus* including *F. colombiensis* as a junior synonym, as well as assorted comments on other species of the *F. acus* group in Colombia.

### *Farlowella acus* (Kner, 1853)

*Acestra acus* Kner, 1853.

*Loricaria scolopacina* De Filippi, 1853:166.

*Farlowella scolopacina* (De Filippi, 1853). In compendia of Loricariid species: Isbrücker (1980:101), Isbrücker (1981:89), Burgess (1989:441).

*Farlowella acus* (Kner, 1853). In compendia of Loricariid or Siluriform species: Eigenmann & Eigenmann (1889:34), Isbrücker (1980:96), Burgess (1989:440), Isbrücker (2001:25,27), Isbrücker (2002:15), Ferraris in Reis et al. (2003:332), Ferraris (2007:237). In taxonomic revisions or species descriptions in Farlowella: Retzer & Page (1997:59), Ballen & Mojica (2014:140), Ballen et al. (2016a:6), Ballen et al. (2016b:331). In geographic faunal checklists: DoNascimiento et al. (2017:76), Urbano-Bonilla et al. (2018:74).

*Farlowella colombiensis* Retzer & Page, 1997. In compendia of Loricariid or Siluriform species: Isbrücker (2001:27), Ferraris in Reis et al. (2003:332), Ferraris (2007:237). In taxonomic revisions or species descriptions in *Farlowella*: Ballen & Mojica (2014:140), Ballen et al. (2016b:331). In geographic faunal checklists: DoNascimiento et al. (2017:76)

Holotype. NMW 47795, 162.64 mm SL, male, Venezuela, Caracas, 8th of August of 1850.

Other types. MZUT 22, not measured, Caracas, Venezuela, holotype of *Loricaria scolopacina* de Filippi, 1853. CAS-SU (ICH) 123733, 132.72 mm SL, male, Departamento del meta, Guaicaramo, río Upia drainage, January of 1928, Holotype of *Farlowella colombiensis* Retzer & Page, 1997.

#### Redescription

Largest specimen, a male, 174.97 mm SL (MPUJ 7321).Body elongate, slender and cylindrical in transversal section. Greatest body depth and width at pelvic-fin insertion. Head slightly depressed, body trunk cylindrical, tail depressed. Dorsal profile slightly concave on lateral view, from snout tip to posterior margin of eyes, relatively straight from that point to dorsal-fin origin, and then straight from last dorsal-fin ray to last caudal-fin plate. Ventral profile relatively straight from snout tip to pelvic-fins insertion, slightly concave from that point to anal-fin origin, and then straight to caudal peduncle. Body completely covered with bony plates except for snout tip, and oval region surrounding urogenital area, small and numerous platelets covering gular region. Snout relatively short, papillae absent. Proportion between snout-mouth length and interorbital distance ≤ 1.86, showing individual variation (Figure 1d-h). Anterior and posterior nares of similar size, dermal flap separating both openings. Eyes lateral, not visible in ventral view. Eyes not set above head surface; iris operculum present. Orbital diameter 0.071–0.098, interorbital diameter 0.182–0.431, and nare-eye distance 0.022–0.075. Sixth infraorbital plate evident. Dorsal surface of head sometimes with longitudinal keel on parieto-supraoccipital bone; compound pterotic ornamented with reticulate pattern of ridges and pits. Mature males with nuptial odontodes on snout, sometimes also on preorbital crest and/or in patches on anterodorsal surface of head (Figure 1d-f). Mouth ovoid, lower lip larger than upper lip; ventral surface covered by wide oval papillae on upper lip and round papillae on lower lip; round papillae decreasing in size from oral aperture to lip margin; lip margin papillose. Few platelets covering dorsal surface of upper lip. Premaxillary with 21 to 34 bicuspid teeth. Dentary with 18 to 33 bicuspid teeth; premaxilla wider than dentary. Buccal papilla present, with papillose surface. Maxillary barbels small and projecting slightly from mouth margin. Flat abdomen covered with two series of plates, complete and continuous, although some individuals show anterior median incomplete series with up to seven plates. Dorsal plates 32-35, dorsomedian plates 12–18, coalescent plates 16–23, ventromedian plates 13–19, and ventral plates 33–38. Dorsal-fin rays i-6, pectoral-fin rays i-6, pelvic-fin rays i-(4-5), caudal-fin rays i-10-i.

#### Colouration

Background coloration light brown, with lateral darker stripes that run from snout to end of dorsal fin, leaving a central light brown area or stripe between them from the origin of dorsal fin and vanish into the snout that smoothly becomes darker, the snout is marbled until its tip. The fins are hyaline with darker segments interleaved with lighter ones (Figure 1d-h). The caudal fin with an almost complete half-moon. Some individuals with dark vermiculations head, sometimes limited to the plates (Figure 1d) or not (Figure 1e).

#### Distribution

*Farlowella acus* is widely distributed in the Orinoco basin, in particular in the Meta and Ariari rivers, departments of Arauca, Casanare, Cundinamarca, and Meta (Figure 3c-d).

#### Remarks

Our results suggest that there are no significant morphological differences between *F. acus* and *F. colombiensis*, which is also the case for the characters suggested by Retzer & Page (1997). This is also the case for the caudal colour pattern, as we did not find a distinctive pattern among species (Figure 1d-h). This is supported by the statistical analyses. The PCA (Figure 2) shows complete overlap between *F. acus* and *F. colombiensis*. Futhermore, characters suggested by the PCA as important among the morphometric variables such as snout-eye and snout-mouth lengths did not show statistical differences (p = 0.7296 and p = 0.796, respectively) when including the body depth at the base of the dorsal fin as covariate for representing body size (Figure 4a-b). Furthermore, the distribution pattern of both species overlaps completely near the type locality of *F. colombiensis* (Figure 3c-d).

**Figure 4.**
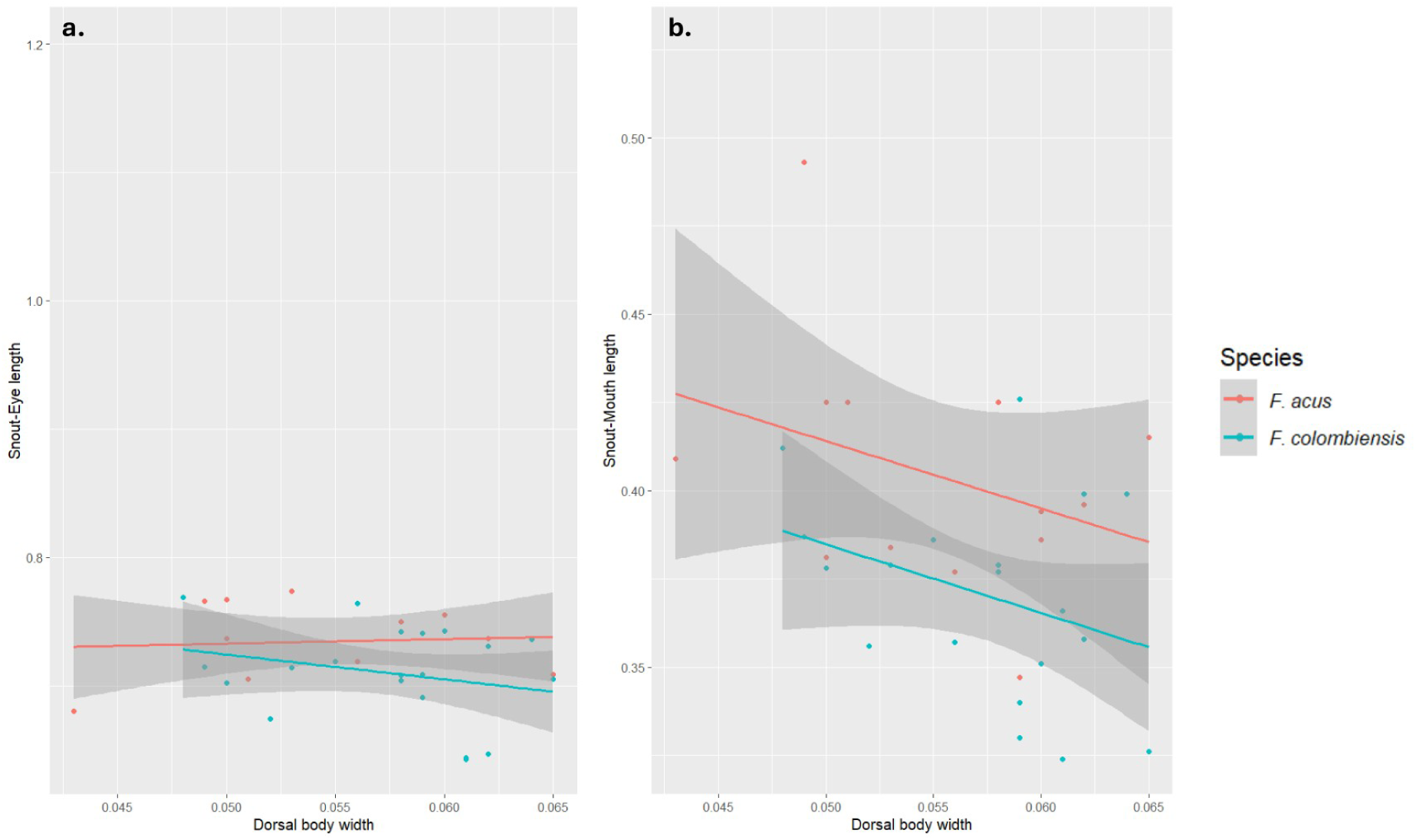
Lack of differences between entre *Farlowella acus* and *Farlowella* “*colombiensis*”; a. snout-eye length, and b. snout-mouth length. The ANCOVA uses the body width at the insertion of the dorsal fin as a covariate.

We note herein for the first time the presence of a bilateral papilla on the anterior margin of the opercular membrane, located anterior to it, and by the plates anterior to the opercular membrane, bearing melanophores (Figure 5). The anatomical nature of such structure is however unclear at the moment, but we observed it in *F. acus*, *F. mitoupibo*, and *F. vittata*. Such structure shows individual variation, and is seen always in a somewhat low frequency (15–23% of the specimens examined for each species). We call it herein the preopercular membrane, not because of any association to the preopercular bone but rather because of its association to the opercular membrane.

**Figure 5.**
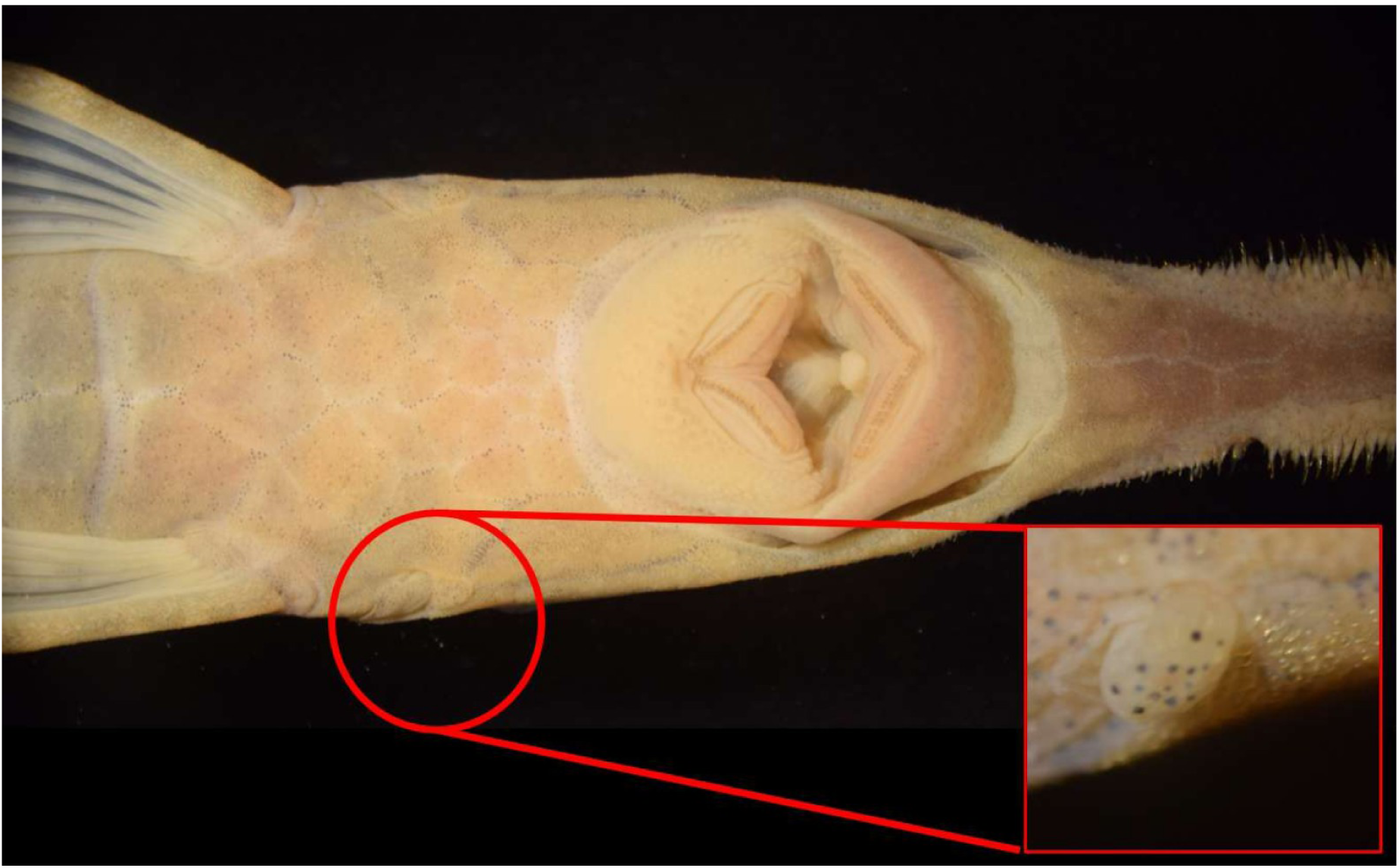
Preopercular papilla in *F. acus*; MPUJ 7320.

### Remarks on *Farlowella mitoupibo*

The species is distributed in tributaries of the Ariari, Guayabero, and Guayuriba rivers of the department of Meta. Herein we record for the first time the presence of the species in the Vaupés river, department of Guaviare (Figure 3e).

### Remarks on *Farlowella vittata*

The morphology of the snout (Figure 1b-c) y morphometric data (Tabla 1) reveal that *F. vitttata* show the longest snout among species of the group. The proportion between snout-mouth length and interorbital distance is larger than in other species except *F. mitoupibo* (≥ 1.88 vs. ≤ 1.86). This shows that such character is useful for separating both species.

*Farlowella vittata* has the widest distribution among species of the *F. acus* group in Colombia, being present in almost all tributaries of the Meta river, in the departments of Arauca, Casanare, Cundinamarca, and Meta. It is also present in the Ariari river, in the departments of Meta and Guaviare (Figure 3b).

### Sexual dimorphism in the *F. acus* group

We found no statistical evidence of sexual dimorphism in morphometric and meristric variables in the *F. acus* group (PCA) (Figure 6). Although we found positive correlation between pectoral-spine length and body size (correlation 0.801), and proportion of snout-mouth length to interorbital distance (correlation −0.818), none of them allowed separation between sexes.

**Figure 6.**
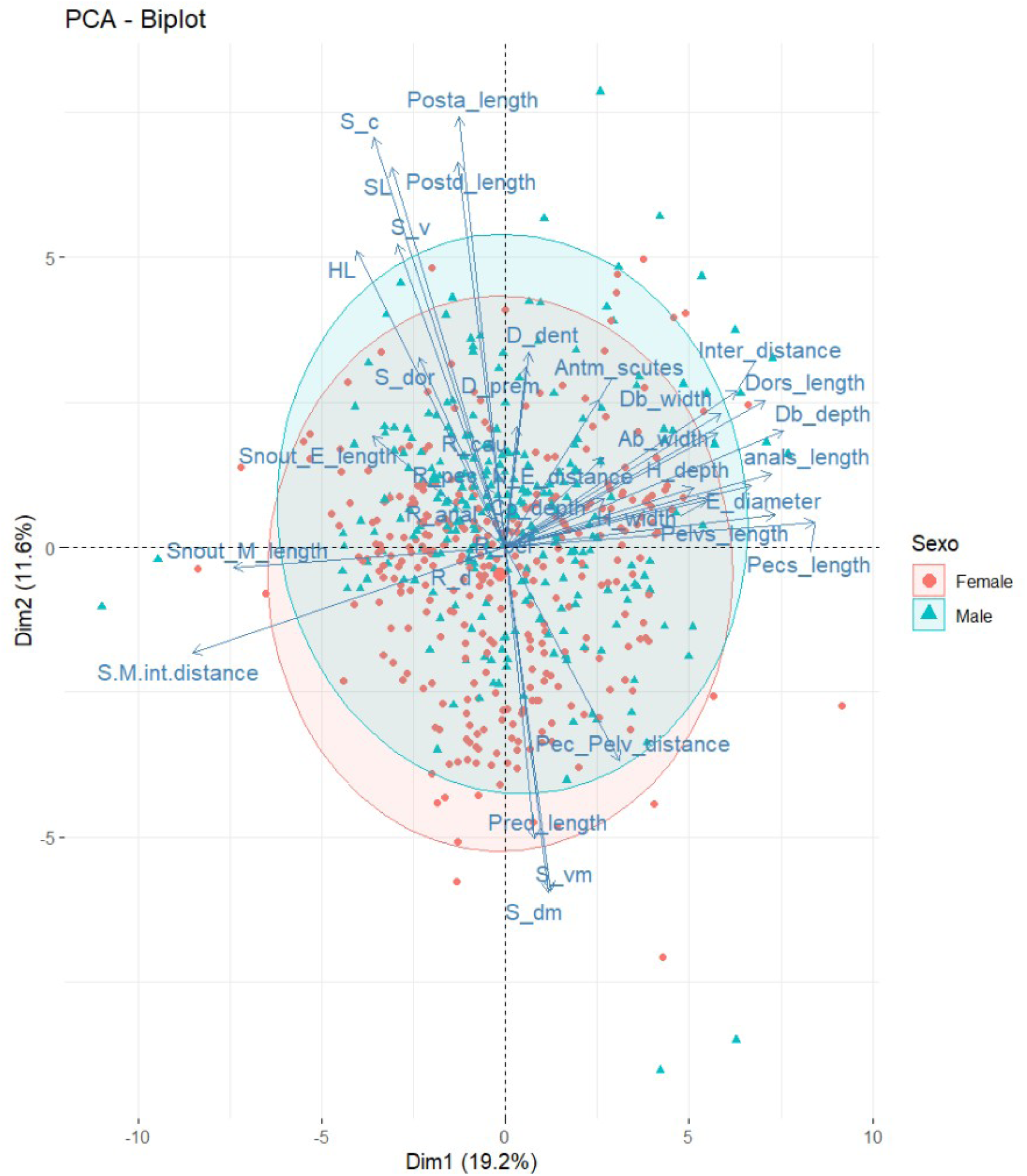
Principal component analysis for exploring possible differences between sexes in the genus *Farlowella*. SL:Standard length, HL:Head length, Db_depth:Dorsal body depth, Db_width:Dorsal body width, Ab_width:Anal body width, Pred_length:Predorsal length, Postd_length:Postdorsal length, Posta_length:Postanal length, Cp_depth:Caudal peduncle depth, Dors_length:Dorsal spine length, Pecs_length:Pectoral spine length, Pelvs_length:Pelvic spine length, Anals_length:Anal spine length, Pec_Pelv_distance:Pectoral-Pelvic distance, Snout_E_length:Snout-eye length, Snout_M_length:Snout-mouth length, E_diameter:Eye diameter, Inter_distance:Interorbital distance, H_width:Head width, H_depth:Head depth, N_E_distance:Narine-eye distance, S-M/int-distance:Snout-mouth / interorbital distance, S_dor:Dorsal plates, S_dm:Dorsomedia-media plates, S_c:Coalecente plates, S_vm:Ventromedia plates, S_v:Ventral plates, R_d:Dorsal rays, R_pec:Pectoral rays, R_pel:Pelvic rays, R_anal:Anal rays, R_cau:Caudal rays, D_prem:Premaxillary teeth, D_dent:Dentary teeth, Antm_scutes:Number of anterior median amdominal scutes.

There is structured variation in the urogenital papilla between mature males and females (Figure 1a). Males have a pointed urogenital papilla, whereas mature females have a swollen urogenital papilla with the aperture proximal to the anus.

## Discussion

### Taxonomy of the *F. acus* species group

Retzer & Page (1997) described *F. colombiensis* using two diagnostic characters: The colour pattern on the caudal fin, and the presence of up to four median ventral abdominal plates in an incomplete series. We show here that there is enough individual variation in both traits to render them unsuitable for separating it from congeners. Furthermore, the incomplete mid-abdominal plate series is also found in both *F. vittata* and *F. acus*. Previous studies recognised the presence of an incomplete mid-abdominal plate series (e.g., Eigenmann & Vance, 1917; Martín Salazar, 1964) with up to two or three anterior plates in *F. acus*. This feature has been used also to identify *F. mitoupibo*, and has been mentioned recently by Dopazo et al. (2023) as one of three possible arrangements of this plate series in *F. wuyjugu*.

The presence of odontodes on the preorbital crest shows individual variation inside a given species (Figure 1g-h); therefore, it is a questionable character for diagnosing species, and possibly for identifying them too. Until now it is unclear whether it also corresponds to sexual or ontogenetic variation, in which case the character may be useful within the limits of its variation. This character has been used in the literature as an aid for identifying *F. acus* (Ballen & Mojica, 2014; Ballen et al., 2016b; Urbano-Bonilla et al., 2018).

Martín-Salazar (1964) suggested the possibility of further splitting *F. acus* in several subspecies, describing *F. acus venezuelensis*. Furthermore, he also noted individual variation in *F. acus*, which we also noted in the morphology of the snout, odontodes, and caudal fin (Figure 1.d-h). That subspecies was eventually erected to the specific level by Retzer & Page (1997). As a consequence of our own findings, it is key to reassess the status of *F. venezuelensis* and *F. martini* which may also happen to be synonyms of *F. acus*, rendering the species widespread and variable. This however is outside the scope of the present work and constitutes future research on this species group.

### Sexual dimorphism in *Farlowella*

This is the first detailed report of sexual dimorphism in the genital papilla in both *Farlowella*, and the subfamily Loricariinae. The same configuration in the genital papilla has been found by Ballen & Vari (2012) in *Ancistrus*, *Cordylancistrus*, *Dolichancistrus*, *Lasiancistrus*, and *Leptancistrus,* by Ballen (2011), Ballen et al., (2016c), Urbano-Bonilla & Ballen (2021), and Meza-Vargas et al. (2022) in *Chaetostoma*, and by Armbruster & Taphorn (2013) in *Neblinichthys.* All of these genera are however outside of the subfamily Loricariinae.

### Key to the species of the *F. acus* species group in Colombia

**1.** Presence of brown margin bordering the ventral plates, median abdominal row discontinuous and incomplete, with 4-6 ventral plates… *Farlowella mitoupibo*. **1’.** Presence of brown margin bordering the ventral plates, median abdominal row either absent or present but continuous… 2
**2.** Snout short and robust (snout-mouth distance to interorbital distance ≤ 1.80)… *Farlowella acus* **2’.** Snout long and slender (snout-mouth distance to interorbital distance > 1.85)… *Farlowella vittata*

## Conclusions

This study contributes towards the understanding of the taxonomy of the species in the *F. acus* species group by highlighting areas of future improvement. We also report for the first time anatomical structures of known (genital papilla) and unknown (preopercular papilla) function, even though it has been thoroughly studied by several authors. We also suggest the possible relationship between snout morphology and geographic distribution. Future research directions include the use of methods from population genetics and phylogenetics in order to tackle the question of the status of wide-ranging species such as *F. acus* and *F. vittata*, which may also shed light on their evolution in space by taking an integrative taxonomic approach

### Specimens examined

#### Farlowella acus

ANSP 130038, 12, 112.62-148.8 mm SL, Venezuela, Carabobo, Río Vigirima, 10°16’44.57"N, 67°57’17.96"W. INHS 60015, 2, 77.57-82.9 mm SL, Venezuela, Carabobo, Lago Valencia. MPUJ 2889, 1, 153.29 mm SL, Colombia, Meta, San Martín, 3° 39’ 36" N, 73° 36’ 36" W. MPUJ 3035, 5, 103.80-159.130 mm SL, Colombia, Meta, Restrepo, Caño Caney, 4° 16’ 48" N, 73° 33’ 0" W. MPUJ 3036, 10, 77.33-196.24 mm SL, Colombia, Meta, Restrepo, Caño Caney, 4° 16’ 48" N, 73° 33’ 0" W. MPUJ 3141, 2, 83.84-117.52 mm SL, Colombia, Meta, Restrepo, Brazuelo del río Caney, 4° 16’ 48" N, 73° 33’ 0" W. MPUJ 3614, 2, 89.190-110.040 mm SL, Colombia, Meta, Acacías, Finca La Fuente, Caño Tres Rotos, 3° 53’ 24" N, 73° 28’ 48" W. MPUJ 3748, 1, 95,500 mm SL, Colombia, Meta, Acacías, Finca La Fuente, Charco La Rubieta, 3° 53’ 24" N, 73° 28’ 24" W. MPUJ 3819, 2, 125.90-144.01 mm SL, Colombia, Meta, Acacías, Finca El Cedral, Canal de riego, 3° 54’ 36" N, 73° 30’ 36" W. MPUJ 3842, 1, 138.99 mm SL, Colombia, Meta, Meta, Acacías, Río Metica (Guamal - Humadea), 0°, 53’ 52,59’’ N, 0°, 28’ 17,89’’ W. MPUJ 6089, 1 119.56 mm SL, Colombia, Casanare, Tauramena, Quebrada Aguablanca, 5° 0’ 36" N, 72° 47’ 24" W. MPUJ 7310, 1, 173.6 mm SL, Colombia, Casanare, Casanare, Tamara, Río Pauto, 5° 50’ 41,3’’N, 72° 6’17,67’’W. MPUJ 7321, 1, 174.97 mm SL, Colombia, Casanare, Casanare, Tamara, Río Pauto, 5° 50’ 41,3’’N, 72° 6’ 17,60’’W. MPUJ 8357, 1, 130.24 mm SL, Colombia, Casanare, Casanare, Tauramena, Río Cusiana, 5° 1’ 28.3" N, 72° 41’ 21.2" W. MPUJ 8382, 1, 130.44 mm SL, Colombia, Cundinamarca, Cundinamarca, Mambita, Río Guavio, 4° 44’ 26.5" N, 73° 18’ 30.8" W. MPUJ 10127, 1, 144.45 mm SL, Colombia, Meta, Meta, Fuente de Oro,, 3° 19’ 28,5"N, 73° 29’ 59,6"W. MPUJ 11158, 1, 128.32 mm SL, Colombia, Arauca, Arauca, Tame,, 6° 30’ 9,3"N, 71° 45’ 55,1"W. MPUJ 12157, 1, 113.94 mm SL, Colombia, Cundinamarca, Cundinamarca, Medina, Río Humea, 4° 31’ 5,23’’ N, 73° 21’ 24,3’’ W. NMW 47795, 1, 162.64 mm SL, Venezuela, Caracas. CAS-SU (ICH) 23733, 2, 101.52-132.72 mm SL, Colombia, Guicaramo, Río Orinoco, 4°44’10.46"N, 73°3’8.67"W. CAS-SU (ICH) 56823, 5, 111.12-137.32 mm SL, Venezuela, Carabobo, Río Torito, 10°2’39.66"N, 68°3’26.01"W. CAS-SU (ICH) 56825, 9, 42.4-50.15 mm SL, Venezuela, Carabobo, Río Noguera, 10° 3’ 20.31" N, 68°4’28.01"W. CAS-SU (ICH) 56851, 1, 125.18 mm SL, Venezuela, Carabobo, Río Guigue, 10°5’1.25"N, 67°47’17.85"W. CAS-SU (ICH) 56852, 3, 118.25-132.37 mm SL, Venezuela, Carabobo, Río Limon, 10°5’1.25"N, 67°47’17.85"W. CAS-SU (ICH) 56854, 1, 126.37 mm SL, Venezuela, Carabobo, Río Guayas, 10°10’37.18"N, 67°55’32.96"W. CAS-SU (ICH) 56855, 4, 117.14-145.48 mm SL, Venezuela, Aragua, Río Aragua, 10°12’41.69"N, 67°22’7.32"W.

#### Farlowella martini

INHS 60099, 2, 141.0-170.2 mm SL, Venezuela, Yaracuy, Río Tupe. INHS 60124, 1, 169.31 mm SL, Venezuela, Carabobo, Río Guarataro.

#### Farlowella mitoupibo

MPUJ 3614, 1, 89.190 mm SL, Colombia, Meta, Acacías, Finca La Fuente, Caño Tres Rotos, 3° 53’ 24" N, 73° 28’ 48" W. MPUJ 8481, 1, 203.470 mm SL, Colombia, Meta, Puerto Lleras, Caño Caribe, 3° 12’ 45.3"N, 73° 28’ 40.1"W. MPUJ 12887, 1, 175.26 mm SL, Colombia, Guaviare, Caño Unilla, 2°11’51.3"N, 72°44’58.7"W. MPUJ 12888, 6, 138.73-157.25 mm SL, Colombia, Guaviare, Caño La Tigra, 2°10’56.7"N, 72°50’15.8"W. MPUJ 13116, 1, 127.57 mm SL, Colombia, Guaviare, Caño Toño, 2°09’49.8"N, 72°50’15.8"W. MPUJ 13117, 3, 126.44-165.10 mm SL, Colombia, Guaviare, Río Itilla (Raudal), 1°59’30.4"N, 72°53’14.6"W. MPUJ 13118, 2, 120.04-129.17 mm SL, Colombia, Guaviare, Río Itilla (Brisas del Itilla), 1°58’00.2"N, 72°50’27.7"W. MPUJ 13121, 1, 110.78 mm SL, Colombia, Guaviare, Caño Potosí, 2°12’18.4"N, 72°38’14.7"W. MPUJ 13205, 1, 120.22 mm SL, Colombia, Guaviare, Caño Bálsamo (Finca el Vergel), 1°57’59.0"N, 72°37’16.5"W.

#### Farlowella venezuelensis

MBUCV-V 11950, 8, 136.33-197.5457 mm SL, Venezuela, Motatan, Río Colorado. MCNG 16971, 5, 61.45-167.4 mm SL, Venezuela, Monagas, Río Cocoyal, 10°7’0"N, 63°41’0"W. USNM 163179, 3, 110.2-160.86 mm SL, Venezuela, Monagas, Río Guarapiche.

#### Farlowella vittata

AMNH 58180, 1, 115.18 mm SL, Venezuela, Caño Mato, 9°9’0’’N, 61°3’0’’ W. ANSP 128728, 9, 119.16-181.58 mm SL, Colombia, Meta, Raucho el Viento, 4°8’32.63"N, 72°39’23.06"W. ANSP 128734, 1, 198.5 mm SL, Colombia, Meta, Cano Emma, 4°5’43"N, 72°24’49"W. ANSP 133319, 1, 98.57 mm SL, Colombia, Meta, Quebrada Venturosa, 4°3’50.35"N, 72°58’53"W. ANSP 134772, 1, 123.44 mm SL, Colombia, Meta, Cano El Chocho, 4°6’45"N, 73°6’40.18"W. ANSP 135726, 7, 98.71-128.21 mm SL, Venezuela, Bolivar, Río Tauca, 7°28’7.39"N, 64°52’58.22"W. ANSP 139618, 1, 132.95 mm SL, Venezuela, Bolivar, Río Mato, 6°47’53.93"N, 65°14’44.72"W. ANSP 159935, 2, 126.72-136.5 mm SL, Venezuela, Bolivar, Cano Caiman, 7°21’16.99"N, 66°15’52.47"W. ANSP 159936, 1, 150.17 mm SL, Venezuela, Bolivar, Río Parguaza, 6°15’38.12"N, 67°7’31.54"W. ANSP 159937, 1, 124.42 mm SL, Venezuela, Bolivar, Río Orocopiche, 8°3’4.8"N, 63°40’3.31"W. ANSP 159938, 1, 161.72 mm SL, Venezuela, Bolivar, Río Caura, 7°36’56.75"N, 64°51’48.96"W. CAS 136513, 2, 90.33-129.98 mm SL, Venezuela, Río Apure. CAS 38145, 1, 89.36-114.17 mm SL, Venezuela, Bolivar, Río Marquarita. CAS 77119, 4, 103.78-127.87 mm SL, Colombia, Meta, Río Meta. FMNH 73440, 1, 131.17 mm SL, Colombia, Meta, Río Guapaya, 3°3’42.48"N, 73°50’26.49"W. FMNH 73441, 1, 154.22 mm SL, Colombia, Meta, Río Guapaya, 3°3’42.48"N, 73°50’26.49"W. INHS 27676, 10, 109.32-224.65 mm SL, Venezuela, Apure, Cañoo Potrerito, 6°24’0’’N, 67°31’0’’W. INHS 27752, 10, 102.50-151.37 mm SL, Venezuela, Barinas, Caño Curito, 7°58’0’’N, 71°1’0’’W. INHS 28302, 8, 113.22-142.52 mm SL, Venezuela, Barinas, Río Apure Río Suripa. INHS 28927, 4, 152.82-192.93 mm SL, Venezuela, Cojedes, Río Tinaco. INHS 59902, 2, 106.60-152.32 mm SL, Venezuela, Portugesa, Río Portugesa - Río Are. INHS 60082, 4, 164.13-173.33 mm SL, Venezuela, Cojedes, Río Cojedes Río Portuguesa. INHS 60145, 7, 102.57-124.90 mm SL, Venezuela, Guarico, Río San Antonio. INHS 60235, 3, 100.05-191.65 mm SL, Venezuela, Cojedes, Río Pao, 9°32’0’’N, 68°7’0’’ W. INHS 60259, 6, 110.89-169.02 mm SL, Venezuela, Carabobo, Río Portuguesa-Río Chirguito. INHS 61248, 1, 146.07, Venezuela, Portuguesa, Río Apure, 5°35’0’’N, 67°42’0’’W. INHS 61279, 7, 134.8-170.35 mm SL, Venezuela, Barinas, Río Santa Barbara, 7°50’0’’N, 71°11’0’’W. INHS 61318, 1, 120.18 mm SL, Venezuela, Barinas, Río Apure Río Canagua. INHS 61555, 10, 150.62-206.1 mm SL, Venezuela, Amazonas, Caño Pozo Azule, 9°9’0’’N, 61°3’0’’W. INHS 62021, 10, 56.36-132.32 mm SL, Venezuela, Guarico, Río Mocapra. INHS 69218, 5, 93.69-117.62 mm SL, Venezuela, Guarico, Río San Jose. INHS 69259, 2, 137-42-156.07 mm SL, Venezuela, Portugesa, Los Manires Creek. INHS 69293, 3, 123.62-137.78 mm SL, Venezuela, Bolivar, Río Suapire. INHS 69304, 2, 141.88-153.44 mm SL, Venezuela, Bolivar, Río Urbani. INHS 69414, 5, 45.49-118.52, mm SL, Venezuela, Guarico, Río San Bartolo. INHS 69470, 2, 101.05-107.36 mm SL, Venezuela, Guarico, Río Guariquito. INHS 69489, 1, 141.39 mm SL, Venezuela, Portugesa, Río Guanare. INHS 69547, 7, 95.51 - 123.46 mm SL, Venezuela, Guarico, Río San Jose. MBUCV-V 11939, 1, 114.45 mm SL, Venezuela, Bolivar, trib., Orinoco. MBUCV-V 11940, 1, 111.78 mm SL, Venezuela, Cojedes-Carab, Río Apure. MBUCV-V 11943, 1, 163.15 mm SL, Venezuela, Apure, Río Apure. MBUCV-V 11949, 1, 115.44 mm SL, Venezuela, Carabobo, Río Tirgua. MBUCV-V 13243, 5, 83.29-94.53 mm SL, Venezuela, Isla Tres Canos. MCNG 10297, 1, 116.64 mm SL, Venezuela, Apure, Cano Maporal, 7°25’19.92"N, 69°35’39.84"W. MCNG 10733, 6, 97.5-147.90 mm SL, Venezuela, Barinas, Río Pedraza, 7°56’9.96"N, 71°3’20.16"W. MCNG 11045, 2, 87.36-111.37 mm SL, Venezuela, Apure, Cano Maporal, 7°28’40.08"N, 69°31’9.84"W. MCNG 11174, 2, 118.92-172.26 mm SL, Venezuela, Bolivar, Cano Garrapata, 6°19’19.92"N, 67°7’0.12"W. MCNG 11202, 4, 80.17-88.57 mm SL, Venezuela, Monagas, Río Uracoa, 8°45’0"N, 62°46’0.12"W. MCNG 11424, 5, 94.72-130.72 mm SL, Venezuela, Guarico, Río Agua Blanca Caño, 8°43’30"N, 66°53’49.92"W. MCNG 11916, 16, 118.09-139.78 mm SL, Venezuela, Tachira, Río Grande, 7°23’9.96"N, 71°57’39.96"W. MCNG 12539, 1, 100.9 mm SL, Venezuela, Amazonas, Río Cantanaipo. MCNG 13684, 2, 114.27-143.19 mm SL, Venezuela, Cojedes, Río Pao, 9°34’19.92"N, 68°10’40.08"W. MCNG 14287, 1, 105.78 mm SL, Venezuela, Cojedes, Río Orupe, 9°40’40.08"N, 68°30’29.88"W. MCNG 15205, 2, 88.04-103.08 mm SL, Venezuela, Bolivar, Suapire. MCNG 15938, 2, 93.72-108.63 mm SL, Venezuela, Monagas, Río Yabo, 8°54’0"N, 62°50’0"W. MCNG 15963, 10, 95.79-163.69 mm SL, Venezuela, Bolivar, Río Chaviripa, 7°8’0"N, 66°30’0"W. MCNG 16803, 3, 136.14-93.35 mm SL, Venezuela, Guarico, Caño el Sur de El Pilar, 10°30’0"N, 63°7’0.12"W. MCNG 16828, 4, 80.88-93.38 mm SL, Venezuela, Monagas, Cano Agua Clara, 8°57’0"N, 62°48’0"W. MCNG 16861, 3, 72.3-79.06 mm SL, Venezuela, Monagas, Q. del Medio, 8°52’0.12"N, 63°10’0.12"W. MCNG 16924, 5, 73.48-118.24 mm SL, Venezuela, Monagas, Monagas, 9°15’0"N, 62°38’0"W. MCNG 17024, 5, 93.81-15.27 mm SL, Venezuela, Sucre, Río Pilar, 10°32’0"N, 63°8’0"W. MCNG 17046, 7, 88.1-146.7 mm SL, Venezuela, Sucre, Cano Juan Antonio, 10°23’0"N, 63°22’0"W. MCNG 17274, 7, 99.51-142.47 mm SL, Venezuela, Anzoategui, Río Tigre, 8°57’29.88"N, 63°22’0.12"W. MCNG 17741, 10, 121.16-200.30 mm SL, Venezuela, Bolivar, Cano Agua Mena, 6°22’0’’N, 64°56’0’’W. MCNG 20842, 1, 101.64 mm SL, Venezuela, Bolivar, Río Nichare, 6°13’46.92"N, 64°56’18.96"W. MCNG 20879, 2, 100.12-162.12 mm SL, Venezuela, Bolivar, Río Nichare, 6°12’34.92"N, 64°56’35.16"W. MCNG 22007, 1, 142.5 mm SL, Venezuela, Bolivar, Cano Icutu, 5°52’59.88"N, 64°51’0"W. MCNG 22402, 3, 160.06-197.00 mm SL, Venezuela, Bolivar, Río Caura, 5°52’59.88"N, 64°51’0"W. MCNG 22522, 1, 179.32 mm SL, Venezuela, Bolivar, Río Nichare, 6°4’54.84"N, 64°55’0.12"W. MCNG 22868, 1, 167.45 mm SL, Venezuela, Bolivar, Río Nichare, 6°5’15"N, 64°56’3.12"W. MCNG 23021, 4, 79.42-104.00 mm SL, Venezuela, Bolivar, Río Nichare, 6°22’1.92"N, 64°58’0.84"W. MCNG 23276, 6, 89.67-117.92 mm SL, Venezuela, Bolivar, Río Nichare. MCNG 23515, 1, 152.07 mm SL, Venezuela, Tachira, Río Camburito, 7°41’20.04"N, 71°32’9.96"W. MCNG 23827, 2, 109.52-121.73 mm SL, Venezuela, Bolivar, Río Chivaripa, 7°10’0"N, 66°30’0"W. MCNG 23938, 3, 109.82-146.47 mm SL, Venezuela, Amazonas, Río Ocamo, 3°10’59.88"N, 64°31’0.12"W. MCNG 6223, 2, 95.92-129.96 mm SL, Venezuela, Apure, Río Riecito, 6°25’0.12"N, 67°32’0.96"W. MCNG 6597, 5, 121.62-142.25 mm SL, Venezuela, Barinas, Río Apure, 7°42’29.88"N, 71°17’49.92"W. MCNG 6724, 2, 95.91-149.52 mm SL, Venezuela, Bolivar, Cano Trapichote, 6°34’40.08"N, 66°49’30"W. MCNG 6745, 3, 129.62-151.5 mm SL, Venezuela, Bolivar, Río Simonero, 6°37’0’’N, 67°2’0’’W. MCNG 6794, 10, 133.6-181.14 mm SL, Venezuela, Cojedes, Río Portuguesa, 9°39’39.96"N, 68°44’49.92"W. MCNG 7397, 3, 143.16-165.65 mm SL, Venezuela, Barinas, Río Suripa, 7°49’30"N, 70°45’29.88"W. MCNG 7442, 5, 96.47-151.67 mm SL, Venezuela, Barinas, Río Santo Domingo, 8°37’30"N, 70°10’59.88"W. MHNG 2532.28, 2, 108.98-118.98 mm SL, Colombia, Acacias, Río Meta. CAS-SU (ICH) 56824, 1, 140.76 mm SL, Venezuela, Lara, Río Tocuyo, 9°50’4.2"N, 70°13’2.68"W. CAS-SU (ICH) 56826, 1, 134.09 mm SL, Venezuela, Carabobo, Río Bejuma, 10°10’37.4"N, 68°16’0.15"W. UF 26020, 5, 114.78-16942 mm SL, Colombia, Meta, Río Yucao, 4°18’52.84"N, 72°7’42.56"W. UF 26181, 6, 117.6-130.86 mm SL, Colombia, Meta, Río Meta, 4°7’27.44"N, 73°28’56.49"W. UF 33224, 3, 118.88-187.78 mm SL, Colombia, Meta, Río Meta, 4°3’42.71"N, 73°12’24.92"W. UF 33290, 6, 125.22-162.09 mm SL, Colombia, Meta, Río Orotoy, 3°53’12.65"N, 73°23’39.85"W. UF 33331, 2, 139.18-148.25 mm SL, Colombia, Meta, Río Humea, 4°22’59.5"N, 73°11’33.64"W. UF 33419, 5, 103.93-137.38 mm SL, Colombia, Meta, Río Guaviare, 4°17’7.37"N, 73°29’8.84"W. UF 33481, 5, 123.28-165.00 mm SL, Colombia, Meta, Río Yucao, 4°18’52.84"N, 72°7’42.56"W. UF 33512, 6, 178.94-214.37 mm SL, Colombia, Meta, Río Meta, 4°22’58.01"N, 73°5’28.96"W. UF 33600, 2, 136.83-177.33 mm SL, Colombia, Meta, Río Meta, 4°3’28.6"N, 73°0’6.08"W. USNM 265669, 1, 94.8 mm SL, Venezuela, Bolivar, Río Orinoco, 8°22’12"N, 62°42’0"W. MPUJ 1875, 1, 88.97 mm SL, Colombia, Meta, Acacías, Caño San Luis, 3° 54’ 0" N, 73° 35’ 24" W. MPUJ 2734, 3, 90.19-169.76 mm SL, Colombia, Meta, San Martín, Caño Camoa, 3° 39’ 36" N, 73° 36’ 36" W. MPUJ 2782, 2, 141.12-152.56 mm SL, Colombia, Meta, San Martín, Río Caño Camoa, Finca Tocancipá 3° 39’ 36" N, 73° 36’ 36" W. MPUJ 2787, 2, 177.46-181.61 mm SL, Colombia, Meta, San Martín, Caño Camoa, 3° 39’ 36" N, 73° 36’ 36" W. MPUJ 2834, 1, 180.57 mm SL, Colombia, Meta, San Martín, Río Ariari, 3° 41’ 24" N, 73° 46’ 12" W. MPUJ 2839, 16, 120,78-169.01 mm SL, Colombia, Meta, San Martín, Río Camoa, 3° 39’ 36" N, 73° 36’ 36" W. MPUJ 2924, 14, 126.36-149.03 mm SL, Colombia, Meta, San Martín, Río Camoa, 3° 39’ 36" N, 73° 36’ 36" W. MPUJ 2925, 2, 133.75-149.51 mm SL, Colombia, Meta, San Martín, Río Camoa, 3° 39’ 36" N, 73° 36’ 36" W. MPUJ 2941, 1, 174.03 mm SL, Colombia, Meta, San Martín, Río Camoa, 3° 39’ 36" N, 73° 36’ 36" W. MPUJ 3035, 14, 103.80-159.130 mm SL, Colombia, Meta, Restrepo, Caño Caney, 4° 16’ 48" N, 73° 33’ 0" W. MPUJ 3036, 55, 77.33-126.94 mm SL, Colombia, Meta, Restrepo, Caño Caney, 4° 16’ 48" N, 73° 33’ 0" W. MPUJ 3118, 1, 101.94 mm SL, Colombia, Meta, Restrepo, Brazuelo del río Caney, 4° 16’ 48" N, 73° 33’ 0" W. MPUJ 3120, 3, 89.61-91.40, Colombia, Meta, Restrepo, Brazuelo del río Caney, 4° 16’ 48" N, 73° 33’ 0" W. MPUJ 3122, 3, 114.62-122.51 mm SL, Colombia, Meta, Restrepo, Río Caney, 4° 16’ 48" N, 73° 33’ 0" W. MPUJ 3124, 8, 121.29-141.54 mm SL, Colombia, Meta, Restrepo, Brazuelo del río Caney, 4° 16’ 48" N, 73° 33’ 0" W. MPUJ 3126, 5, 102.04-112.17 mm SL, Colombia, Meta, Restrepo, Brazuelo del río Caney, 4° 16’ 48" N, 73° 33’ 0" W. MPUJ 3136, 3, 107.94-109.17 mm SL, Colombia, Meta, Restrepo, Brazuelo del río Caney, 4° 16’ 48" N, 73° 33’ 0" W. MPUJ 3137, 15, 93.45-127.77 mm SL, Colombia, Meta, Restrepo, Brazuelo del río Caney, 4° 16’ 48" N, 73° 33’ 0" W. MPUJ 3138, 1, 109.08 mm SL, Colombia, Meta, Restrepo, Brazuelo del río Caney, 4° 16’ 48" N, 73° 33’ 0" W. MPUJ 3139, 3, 129.63-148.78 mm SL, Colombia, Meta, Restrepo, Brazuelo del río Caney, 4° 16’ 48" N, 73° 33’ 0" W. MPUJ 3141, 7, 83.84-117.52 mm SL, Colombia, Meta, Restrepo, Brazuelo del río Caney, 4° 16’ 48" N, 73° 33’ 0" W. MPUJ 3172, 72, 100.47-138.65 mm SL, Colombia, Meta, Restrepo, Río Guatiquía, 4° 17’ 24" N, 73° 34’ 12" W. MPUJ 3173, 1, 131.08 mm SL, Colombia, Meta, Restrepo, Brazuelo del río Caney, 4° 16’ 48" N, 73° 33’ 0" W. MPUJ 3174, 8, 94.94-16.07 mm SL, Colombia, Meta, Restrepo, Brazuelo del río Caney, 4° 16’ 48" N, 73° 33’ 0" W. MPUJ 3181, 4, 114.12-139.04 mm SL, Colombia, Meta, Restrepo, Río Guatiquía, 4° 17’ 24" N, 73° 34’ 12" W. MPUJ 3230, 5, 113.84-120.20 mm SL, Colombia, Meta, Restrepo, Brazuelo del río Caney, 4° 16’ 48" N, 73° 33’ 0" W. MPUJ 3818, 1, 146.82 mm SL, Colombia, Meta, Acacías, Finca El Cedral, Canal de riego, 3° 54’ 0" N, 73° 30’ 36" W. MPUJ 3819, 7, 125.90-144.01 mm SL, Colombia, Meta, Acacías, Finca El Cedral, Canal de riego, 3° 54’ 36" N, 73° 30’ 36" W. MPUJ 3950, 1, 85.15, Colombia, Meta, Acacías, Finca El Cedral, Canal de riego, 3° 54’ 36" N, 73° 30’ 36" W. MPUJ 4152, 1, 107.93 mm SL, Colombia, Meta, Acacías, 3° 53’ 24" N, 73° 27’ 36" W. MPUJ 4454, 2, 154.63-160.31 mm SL, Colombia, Vichada, Puerto Carreño, 6° 0’ 0" N, 67° 25’ 12" W. MPUJ 5801, 1, 125.52mm SL, Colombia, Meta, Puerto López, 4° 16’ 12" N, 72° 36’ 0" W. MPUJ 6050, 1, 119.58 mm SL, Colombia, Casanare, Tauramena, Caño Seco, 4° 58’ 48" N, 72° 44’ 24" W. MPUJ 6067, 3, 96.97-120.52 mm SL, Colombia, Casanare, Tauramena, Quebrada Aguablanca, 5° 0’ 36" N, 72° 47’ 24" W. MPUJ 6083, 1, 157.03 mm SL, Colombia, Casanare, Tauramena, Quebrada Aguablanca, 5° 0’ 36" N, 72° 47’ 24" W. MPUJ 6089, 4, 150.87-110.68 mm SL, Colombia, Casanare, Tauramena, Quebrada Aguablanca, 5° 0’ 36" N, 72° 47’ 24" W. MPUJ 6095, 1, 115.87 mm SL, Colombia, Casanare, Tauramena, Caño Seco, 4° 58’ 48" N, 72° 44’ 24" W. MPUJ 6121, 2, 100.39-149.85 mm SL, Colombia, Casanare, Tauramena, Entre caño Aguablanca y Caño Seco, 4° 58’ 48" N, 72° 44’ 24" W. MPUJ 6127, 3, 96.67-108.39 mm SL, Colombia, Casanare, Tauramena, Quebrada Aguablanca, 5° 0’ 36" N, 72° 47’ 24" W. MPUJ 6700, 1, 107.83 mm SL, Colombia, Casanare, Tauramena, Río Chitamena, 4° 55’ 12" N, 72° 40’ 12" W. MPUJ 6741, 3, 127.80-152.19mm SL, Colombia, Casanare, Tauramena, Orinoquia, Río Caja, 5° 1’ 12" N, 72° 42’ 0" W. MPUJ 6818, 1, 139.32 mm SL, Colombia, Casanare, Tauramena, Orinoquia, Río Chitamena, 4° 54’ 36" N, 72° 31’ 48" W. MPUJ 6854, 1, 91.92 mm SL, Colombia, Casanare, Aguazul, Río Cusiana, 5° 1’ 12" N, 72° 41’ 24" W. MPUJ 6901, 1, 162.72, Colombia, Casanare, Tauramena, Río Cusiana, 5° 0’ 5" N, 72° 41’ 30" W. MPUJ 6915, 1, 114.11 mm SL, Colombia, Casanare, Tauramena, Río chitamena, 4° 55’ 12" N, 72° 40’ 48" W. MPUJ 6938, 2, 121.61-167.35 mm SL, Colombia, Casanare, Aguazul, Desembocadura quebrada Upamena, 5° 0’ 8" N, 72° 41’ 24" W. MPUJ 7118, 3, 99.88-129.05 mm SL, Colombia, Casanare, Tauramena, Río Chitamena, 4° 55’ 48" N, 72° 40’ 12" W. MPUJ 7184, 2, 140.13-140.98 mm SL, Colombia, Casanare, Tauramena, Río Upamena, 5° 1’ 48"N, 72° 41’ 24"W MPUJ 7212, 2, 155.67-179.86 mm SL, Colombia, Casanare, Tauramena, Río Chitamena, 4° 55’ 48"N, 72° 40’ 48"W. MPUJ 7307, 2, 135.44-142.72 mm SL, Colombia, Arauca, Tame, Río Tocoragua, 6° 16’ 12"N, 71° 51’ 36"W. MPUJ 7309, 1, 118.20 mm SL, Colombia, Casanare, Tamara, Quebrada Bayagua, 5° 48’ 0"N, 72° 6’ 36"W. MPUJ 7312, 7, 106.34-146.09 mm SL, Colombia, Arauca, Tame, Caño Caribabare 6° 16’ 48"N, 71° 46’ 12"W. MPUJ 7313, 7, 114.06-152.00 mm SL, Colombia, Arauca, Tame, Caño Puna Puna, 6° 19’ 48"N, 71° 45’ 0"W. MPUJ 7314, 3, 111.19-144.14 mm SL, Colombia, Arauca, Tame, Río Purare, 6° 15’ 36"N, 71° 51’ 36"W. MPUJ 7315, 2, 107.81-137.53 mm SL, Colombia, Arauca, Tame, Río Cravo Norte, 6° 30’ 0"N, 71° 46’ 12"W. MPUJ 7316, 1, 121.57 mm SL, Colombia, Casanare, Tamara, Quebrada La Llorona, 5° 48’ 0"N, 72° 12’ 36"W. MPUJ 7319, 1, 110.27 mm SL, Colombia, Arauca, Tame, Caño Gualabao, 6° 28’ 12"N, 71° 44’ 24"W. MPUJ 7323, 1, 116.73 mm SL, Colombia, Casanare, San Luis de Palenque, Caño Guanapalo, 5° 30’ 36"N, 71° 57’ 0"W MPUJ 7326, 1, 101.77 mm SL, Colombia, Casanare, Tamara, Quebrada La Pone, 5° 45’ 36"N, 72° 6’ 36"W. MPUJ 7327, 1, 100.46 mm SL, Colombia, Casanare, Tamara, Quebrada Bayagua, 5° 43’ 48"N, 72° 7’ 12"W. MPUJ 7328, 7, 121.45-141.30 mm SL, Colombia,Arauca,Tame,Caño Puna Puna, 6° 19’ 48"N, 71° 46’ 12"W. MPUJ 7329, 1, 150.67 mm SL, Colombia, Arauca, Tame, Caño Puna Puna, 6° 19’ 48"N, 71° 46’ 12"W. MPUJ 7374, 1, 116.26 mm SL, Colombia, Casanare, Tamara, Quebrada La Zuquía, 5° 43’ 12"N, 72° 7’ 48"W. MPUJ 7420, 1, 142.77, Colombia, Casanare, Tauramena, Río Caja, 5° 00’ 36"N, 72° 41’ 24"W. MPUJ 7467, 1, 91.07 mm SL, Colombia, Casanare, Tauramena, Río Chitamena, 4° 55’ 48"N, 72° 40’ 12"W. MPUJ 7498, 7, 104.33-1540.01 mm SL, Colombia, Casanare, Tauramena, Quebrada Iglesiera, 5° 02’ 59"N, 72° 47’ 24"W. MPUJ 7546, 1, 90.93 mm SL, Colombia, Casanare, Tauramena, Río Cusiana, 5° 00’ 36"N, 72° 41’ 24"W. MPUJ 7559, 1, 99.85 mm SL, Colombia, Casanare, Tauramena, Río Caja, 5° 00’ 36"N, 72° 41’ 24"W. MPUJ 7561, 3, 92.32-125.15 mm SL, Colombia, Casanare, Tauramena, Río Caja, 5° 01’ 12"N, 72° 42’ 00"W. MPUJ 7579, 1, 137.60 mm SL, Colombia, Casanare, Tauramena, Río Chitamena, 4° 55’ 48"N, 72° 40’ 12"W. MPUJ 7617, 6, 117.57-154.66 mm SL, Colombia, Casanare, Tauramena, Quebrada Iglesiera, 5° 02’ 59"N, 72° 47’ 24"W. MPUJ 7901, 2, 108.28-143.05 mm SL, Colombia, Casanare, Tauramena, Río Caja, 5° 01’ 12"N, 72° 41’ 24"W. MPUJ 7921, 1, 172.69 mm SL, Colombia, Casanare, Tauramena, Río Upanema, 5° 00’ 36"N, 72° 41’ 24"W. MPUJ 8312, 1, 121.51 mm SL, Colombia, Casanare, Tauramena, Quebrada Tauramenera, 5° 00’ 36"N, 72° 45’ 00"W. MPUJ 8325, 1, 136.37 mm SL, Colombia, Casanare, Tauramena, Río Cusiana, 5° 01’ 12"N, 72° 41’ 24"W. MPUJ 8334, 1, 180.23 mm SL, Colombia, Casanare, Tauramena, Río Cusiana, 5° 01’ 12"N, 72° 41’ 24"W. MPUJ 8349, 4, 102.26-125.70 mm SL, Colombia, Casanare, Tauramena, Río Chitamena, 4° 55’ 48"N, 72° 40’ 12"W. MPUJ 9283, 1, 120.08 mm SL, Colombia, Casanare, Tamara, Quebrada La Ceiba, 5° 47’ 24"N, 72° 12’ 00"W. MPUJ 9285, 1, 107.62, mm SL, Colombia, Arauca, Tame, Caño La Florida, 6° 16’ 00"N, 71° 51’ 00"W. MPUJ 9286, 1, 95.34 mm SL, Colombia, Casanare, San Luis de Palenque, Caño Guanapalo, 5° 30’ 36"N, 71° 57’ 00"W. MPUJ 9464, 3, 100.62-144.97 mm SL, Colombia, Casanare, Tauramena, Río Upamena, 5° 01’ 12"N, 72° 41’ 24"W. MPUJ 9488, 1, 145.28 mm SL, Colombia, Casanare, Tauramena, Río Chitamena, 4° 55’ 48"N, 72° 40’ 12"W. MPUJ 9489, 1, 142.28 mm SL, Colombia, Casanare, Tauramena, Río Chitamena, 4° 55’ 48"N, 72° 40’ 12"W. MPUJ 9913, 1, 167.09 mm SL, Colombia, Meta, Vista Hermosa, Río Sardinata, 3° 01’ 12"N, 73° 50’ 24"W. MPUJ 9915, 1, 169.53 mm SL, Colombia, Meta, Macarena, Caño Canoas, 2° 28’ 12"N, 73° 44’ 24"W. MPUJ 9916, 1, 136.53 mm SL, Colombia, Meta, Macarena, Caño Canoas, 2° 28’ 12"N, 73° 44’ 24"W. MPUJ 9919, 5, 130.90-167.01 mm SL, Colombia, Meta, Vista Hermosa, Caño Guapaya, 3° 02’ 24"N, 73° 49’ 48"W. MPUJ 9922, 1, 102.69 mm SL, Colombia, Meta, Vista Hermosa, Caño Blanco, 3° 04’ 48"N, 73° 47’ 24"W. MPUJ 10046, 1, 199.30 mm SL, Colombia, Meta, Mapiripán, Caño La División, 3° 07’ 12"N, 72° 32’ 24"W. MPUJ 10122, 6, 135.88-180.08 mm SL, Colombia,Meta, Caño Irique, 3° 24’ 36"N, 73° 35’ 24"W. MPUJ 10123, 2, 131.56-150.58 mm SL, Colombia, Meta, Puerto Lleras, Caño Abrote, 3° 18’ 00"N, 73° 27’ 00"W. MPUJ 10124, 1, 127.87 mm SL, Colombia, Meta, Caño Guanayas, 3° 22’ 48"N, 73° 40’ 12"W. MPUJ 10125, 1, 123.47, mm SL, Colombia, Meta, San Juan de Arama, Caño Uricacha, 3° 18’ 24"N, 73° 40’ 48"W. MPUJ 10126, 1, 133.16 mm SL, Colombia, Meta, Fuente de Oro, Caño Upín, 3° 21’ 00"N, 73° 40’ 48"W. MPUJ 10129, 2, 113.38-115.57 mm SL, Colombia, Meta, Granada, Caño La Cubillera, 3° 15’ 00"N, 73° 38’ 24"W. MPUJ 10132, 1, 120.42 mm SL, Colombia, Meta, Río Ariari, 3° 44’ 24"N, 73° 43’ 48"W. MPUJ 10135, 1, 111.34 mm SL, Colombia, Meta, Fuente de Oro, Río Ariari, 3° 39’ 36"N, 73° 37’ 48"W. MPUJ 10902, 1, 100.38 mm SL, Colombia, Meta, Vista Hermosa, Caño Guapaya, 3° 02’ 24"N, 73° 49’ 48"W. MPUJ 10915, 1, 115.88 mm SL, Colombia, Meta, Acacías, Caño Agua Caliente y Fría, 3° 30’ 00"N, 73° 43’ 24"W. MPUJ 11564, 2, 149.26-197.22 mm SL, Colombia, Guaviare, San José del Guaviare, Caño Piedra, 2° 21’ 00"N, 72° 49’ 48"W.

## Acknowledgments

GAB was funded through a doctoral scholarship, an internship abroad, and a postdoctoral fellowship by FAPESP (processes #2014/11558-5, #2016/02253-1, and #2023/07838-1). We thank the Museo Javeriano de Historia Natural “Lorenzo Uribe, S.J.” for allowing access to specimens under their care. We thank the comments by T. P. Carvalho during the early phase of assessment of the undergraduate thesis of OEM-O which helped us improve the quality of this work. GAB thanks Alex Urbano-Bonilla, Alejandro Londoño-Burbano, and Manuela Dopazo for discussions on loricariines through the years.

